# Human and bats genome robustness under COSMIC mutational signatures

**DOI:** 10.1101/2024.09.05.611453

**Authors:** Joon-Hyun Song, Ying Zeng, Liliana M. Dávalos, Thomas MacCarthy, Mani Larijani, Mehdi Damaghi

## Abstract

Carcinogenesis is an evolutionary process, and mutations can fix the selected phenotypes in selective microenvironments. Both normal and neoplastic cells are robust to the mutational stressors in the microenvironment to the extent that secure their fitness. To test the robustness of genes under a range of mutagens, we developed a sequential mutation simulator, Sinabro, to simulate single base substitution under a given mutational process. Then, we developed a pipeline to measure the robustness of genes and cells under those mutagenesis processes. We discovered significant human genome robustness to the APOBEC mutational signature SBS2, which is associated with viral defense mechanisms and is implicated in cancer. Robustness evaluations across over 70,000 sequences against 41 signatures showed higher resilience under signatures predominantly causing C-to-T (G-to-A) mutations. Principal component analysis indicates the GC content at the codon’s wobble position significantly influences robustness, with increased resilience noted under transition mutations compared to transversions. Then, we tested our results in bats at extremes of the lifespan-to-mass relationship and found the long-lived bat is more robust to APOBEC than the short-lived one. By revealing robustness to APOBEC ranked highest in human (and bats with much more than number of APOBEC) genome, this work bolsters the key potential role of APOBECs in aging and cancer, as well as evolved countermeasures to this innate mutagenic process. It also provides the baseline of the human and bat genome robustness under mutational processes associated with aging and cancer.

**Highlights:**

- Sinabro, the sequential mutation simulator, facilitates measuring the robustness of human protein-coding sequences under all COSMIC mutational signatures.
- Robustness under APOBEC mutational signatures showed the largest mean and standard deviation in the human genome.
- Robustness to mutational signatures analysis reveals the role of APOBECs is complementary to cancer in the evolvability of cancer cells in later stages.
- Principal component analysis indicates that the GC content at the codon’s wobble position significantly influences robustness.
- A long-lived bat (*Myotis myotis*) has higher robustness to APOBECs than a short-lived one (*Molossus molossus*) than humans.

## Introduction

The evolutionary theory of cancer posits that mutations are key drivers of cancer initiation and progression, as mutation accumulation over time can transform normal cells into malignant cancer cells and also fuel their evolution^1,2^. Therefore, the main focus of much research is studying mutations in cancer as the main factor initiating the disease or shaping the cancer’s ability to survive, adapt, and evolve in response to various challenges. However, what is overlooked is how our genome evolves against all these various types of mutations and mutagenic processes^2–5^.

Based on their response to external and internal stressors, biological systems can be categorized into two groups: robust or evolvable. Robustness is a property of a system to resist internal or external perturbations. Robustness of the biological systems can be found in various levels in organisms, from tissues such as body temperature and homeostasis to molecules such as molecular canalization during development^6^. In contrast, evolvability is a property of a system to maintain or even increase heritable phenotypic variability throughout generations to facilitate adaptation^6–10^.

Cancer is a fast-evolving system that follows both Darwinian and non-Darwinian evolution^10,11^. Understanding our body’s normal cell homeostatic state as a system robust to mutations will illuminate the evolutionary history and principles of carcinogenesis. It can also answer the question of why some of us get cancer, and some don’t, or why some of us die from cancer while some do not^12,13^. There have been several studies to understand cancer’s evolutionary trajectories by profiling cancer cell mutations over time^14–17^. Those studies can record the sequence of events, but miss the causes of those events on the whole genome. Recently, a different approach was developed to examine mutations of the whole genome to define distinct mutational patterns called mutational signatures^18^. The latter gives the opportunity to examine the whole genome as one unit. We used these signatures to measure genome robustness to mutational signatures using our pipeline, which induces one mutation at a time and measures the robustness of each gene to each mutation type. Our hypothesis is that the human genome has evolved to be robust to several mutagens, considering our genome a robust system to stressors.

In this study, we redefine and quantitatively assess the robustness of protein-coding sequences in the human genome under various mutational signatures using our Python-based simulation tool, Sinabro, meaning “little by little unknowingly” in Korean, describing the accumulation of mutations. In this context, robustness is defined as the average number of mutations a sequence can withstand before an amino acid change occurs. Sinabro facilitates the simulation of sequential single-base substitutions within a coding sequence until a predefined stop condition—either a maximum number of mutations or amino acid sequence changes—is reached. We discovered our genome has variable robustness across different mutational signatures with the maximum robustness to signature SBS2, that is, the APOBECs signature. Furthermore, the study explores the robustness of over 70,000 coding sequences against 41 known mutational signatures, revealing that sequences are particularly robust under signatures that predominantly cause C-to-T (G-to-A) mutations. Principal component analysis (PCA) of the robustness data suggests the GC content at the wobble position of codons has a positive correlation determining robustness. Notably, the genome shows increased robustness under transition mutations compared to transversions, aligning with our findings that transition mutation preference correlates positively with robustness. Finally, we compared human genome robustness to APOBECs with two bats: one long-lived (*M. myotis*) and the other one short-lived (*M. molossus*), APOBECs than primates with relatively much less cancer incidence and more extended longevity. We found the long-lived bat had higher robustness compared to humans and the short-lived bat. Our research can open a new avenue on how to analyze the mutation and mutagenesis process in carcinogenesis and cancer outcomes, particularly with respect to aging-induced cancer.

## Results

### Robustness analysis in cancer evolution via Sinabro-based sequential mutation simulation

Robustness is the tendency of the system to be invariant against internal and/or external perturbation. Because systems can have invariance in multiple factors, multiple valid definitions of robustness are possible. We reviewed previous definitions of robustness from the perspective of mutational processes and redefined the robustness under mutational signatures (Supplementary Note S1). Briefly, the definition of the robustness of a protein-coding sequence is the average number of mutations that the sequence can tolerate before amino acid change. We developed the Python program Sinabro to simulate sequential mutation. Sinabro simulates single base substitution under a given mutational process until the stop condition is met. Mutational processes can be simply random, a single mutation type, or COSMIC mutational signatures. A stop condition can be set as the maximum number of mutations or amino acid sequence changes. We applied Sinabro to measure the robustness we defined.

The workflow to measure robustness by Sinabro follows the ensuing steps (Figure 1). First, Sinabro takes a protein-coding sequence starting from the start codon (ATG) to one of the stop codons (TAA, TGA, TAG) with an additional one bp nucleotide, each upstream and downstream. Second, Sinabro computes the probability of nucleotide s_i_ at position *i* to be mutated into a nucleotide a_j_, denoted as P(s_i_>a_j_), based on a given SBS mutational signature. Third, a single mutation is selected based on those probabilities, and the simulator checks if the mutation changed the amino acid sequence of a given protein coding sequence. Fourth, the simulator repeats sequential mutation until the mutation changes the amino acid sequences. Finally, the resulting output of one cycle of complete sequential mutations is a sequence of *n* sequences, and the number of mutations *n*-2 is saved for computation of the robustness. The simulator repeats 1,000 cycles of sequential mutations of the original input sequence, and the average number of mutations is measured as the robustness of the input sequence.

**Figure 1.**
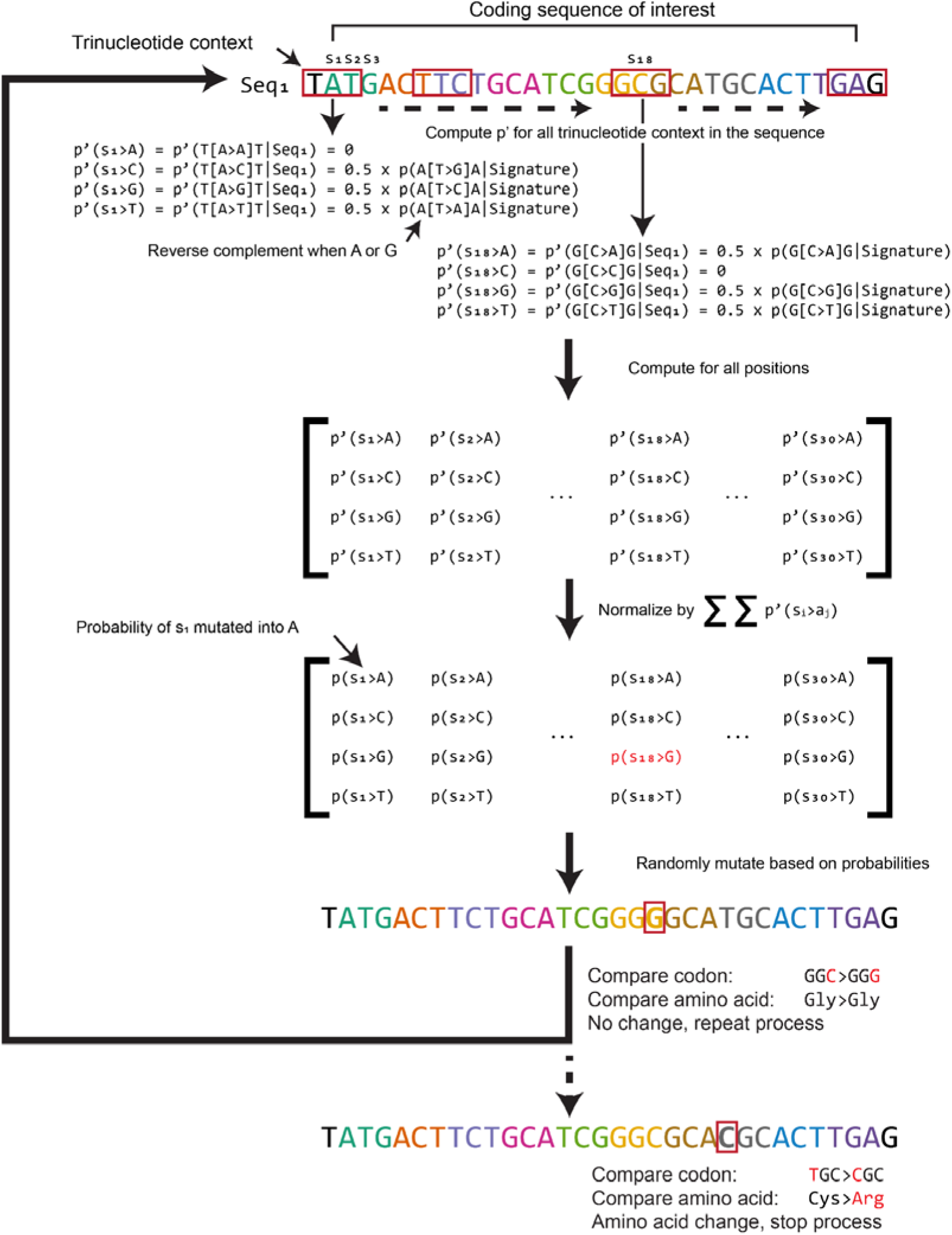
Schematic representation of Sinabro algorithm for sequential mutations under mutational signatures. For a given coding sequence, Sinabro traverses trinucleotide context through the sequence and calls in the percentage of corresponding mutation type based on mutational signature from the pre-computed matrix. The matrix is then normalized to the sum of the matrix to make it a probability matrix. A single base substitution of the sequence is selected based on the probability matrix, and the mutation is compared to the original codon to determine whether it is non-synonymous. The process loops until the mutation is non-synonymous.

We validated Sinabro by simulating a single mutation of each mutational signature for 10,000 random sequences (Figure S1). The average cosine similarity between the original mutational signatures and simulated signatures was 0.974, supporting that Sinabro recapitulates mutational processes given by signatures (Figure S2).

### The human genome is most robust under APOBEC mutational signature SBS2

Applying Sinabro, we computed the robustness against all mutational signatures of more than 70,000 coding sequences of transcripts of the human genome. Surprisingly, APOBEC mutational signatures SBS2 appeared to have both the largest average robustness and variation in robustness (Figures 2a and 2b). APOBEC is a cytidine deaminase that participates in a viral defense mechanism. Recent studies have suspected APOBEC plays a role in cancer formation and intratumor heterogeneity through mutagenesis^19,20^.

**Figure 2.**
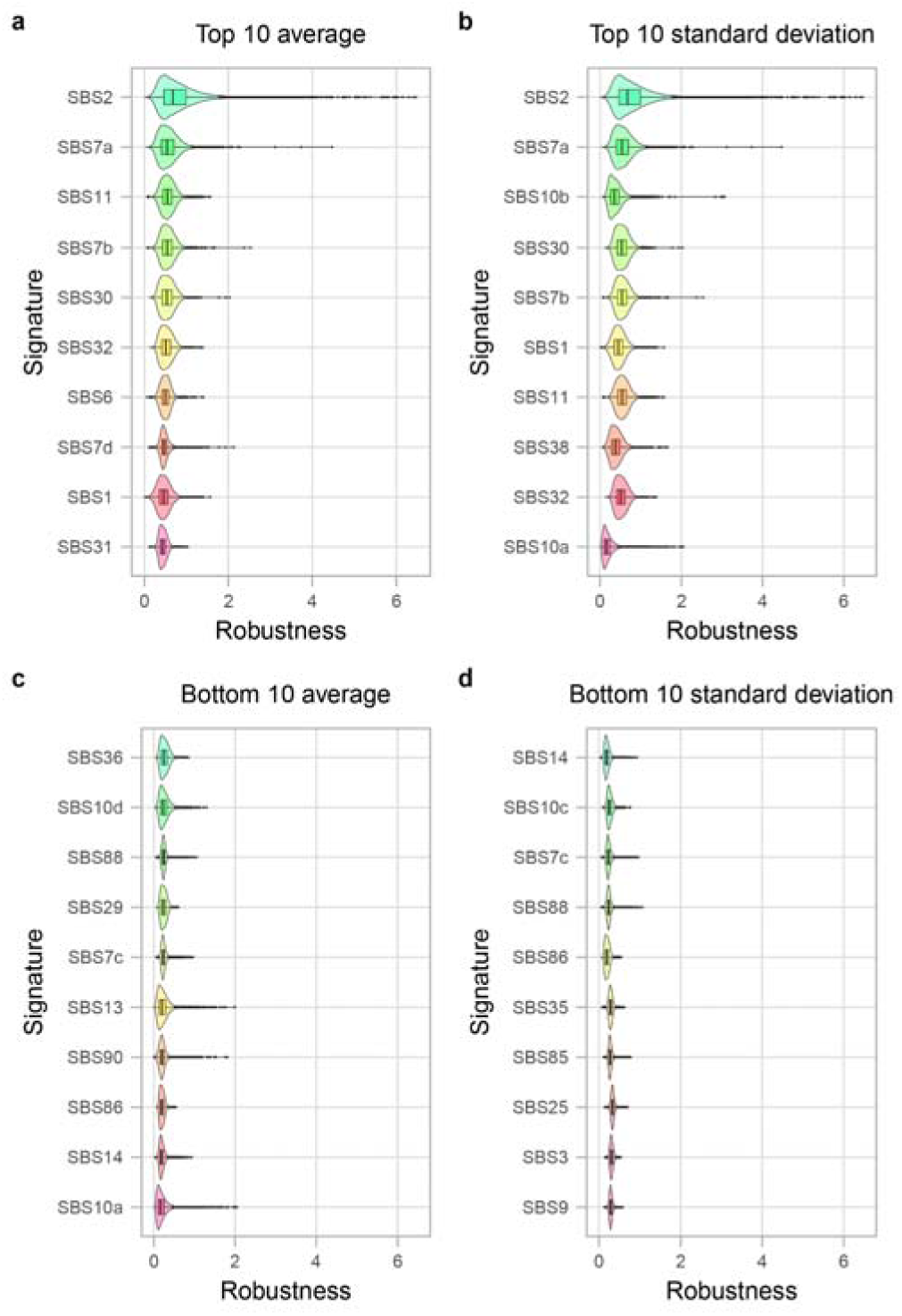
The average and variation of robustness against each mutational signature of the human genome. **a.** The top 10 mutational signatures rank by average. **b.** The top 10 mutational signatures rank by standard deviation. **c.** The bottom 10 mutational signatures rank by average. **d.** The bottom 10 mutational signatures rank by standard deviation.

We first investigated up to the 15th rank based on the average robustness under 41 mutational signatures with known etiology (Figure 2a; Table S1). Robustness under SBS7a, 7b, and 7d are ultraviolet light exposure signatures ranked 2nd, 4th, and 8th. The third mutational signature under which the human genome is robust was SBS11, whose proposed etiology is a temozolomide treatment. SBS11 resembles the mutational signature of alkylating agents^21^. Another mutational signature, SBS31 (platinum chemotherapy treatment), whose patterns are similar to those of alkylating agents, was ranked 10th. There were five mutational signatures associated with a defective DNA repair mechanism; among them, four were an indication of defective DNA mismatch repair (SBS6, SBS15, SBS26, SBS44) and a defective DNA base excision repair due to NTHL1 mutations (SBS30). The robustness under SBS1, a signature of spontaneous deamination of 5-methylcytosine correlated to age that we originally expected to have higher robustness, surprisingly ranked only 9th. The remaining mutational signatures within the top 15 were SBS32 (azathioprine treatment), SBS84 (activity of activation-induced cytidine deaminase), and SBS92 (tobacco smoking), ranked 6th, 11th, and 13th, respectively.

We then further investigated mutational signatures under which the human genome showed low robustness (Figure 2c; Table S1). The bottom 15 mutational signatures were SBS10a (polymerase epsilon exonuclease domain mutations), SBS14 (concurrent polymerase epsilon mutation and defective DNA mismatch repair), SBS86 (unknown chemotherapy treatment), SBS90 (duocarmycin exposure), SBS13 (activity of APOBEC family of cytidine deaminases), SBS7c (ultraviolet light exposure), SBS29 (tobacco chewing), SBS88 (colibactin exposure), SBS10d (defective POLD1 proofreading), SBS36 (defective DNA base excision repair due to MUTYH mutations), SBS10c (defective POLD1 proofreading), SBS24 (aflatoxin exposure), SBS85 (indirect effects of activation-induced cytidine deaminase), SBS18 (damage by reactive oxygen species), and SBS35 (platinum chemotherapy treatment). Nine out of fifteen mutational signatures of the human genome are least robust to induce mainly C-to-A mutations (Figure S3).

### The human genome is robust under mutational processes targeting G/C

To understand common features between mutational signatures under which the human genome is robust, we investigated the targeting preferences of the top 10 mutational signatures (Figure S4). Interestingly, except for SBS7d, all mutational signatures almost exclusively induce C-to-T (G-to-A) mutations. In addition to the average robustness value, the top 10 mutational signatures based on the standard deviation showed a similar pattern that mutational signatures resulting in C-to-T (G-to-A) mutations except for SBS38 and SBS10a which induce C-to-A (G-to-T) mutations (Figure S5). Based on these results, we hypothesize that the GC contents of genes and the GC targeting preference of mutational signatures are crucial to robustness. To examine the hypothesis, we performed a principal component analysis (PCA) on a gene-by-mutational-signature matrix, with each element having a corresponding robustness value. Then, we analyzed principal components by sorting the weight of each mutational signature based on the GC targeting preference of mutational signature (Figure 3a). We found that the weights of the first principal component (PC1) positively correlated to the GC targeting preference of the mutational signature. By contrast, the weights of the second principal component (PC2) negatively correlated to the GC targeting mutational signature except SBS2. The third position of the codon is often called the “wobble” position, which originated from the “wobble” hypothesis by Francis Crick that the 5’ base of the anticodon can form non-standard pairing by “wobble” movement. This makes mutations at the “wobble” position less consequential than those at other positions. Hence, we analyzed the GC contents of the wobble position of the genes on the PC1-PC2 plot (Figure 3c). The result showed that the GC content of the codon’s third position of a gene is positively correlated with PC1 and negatively correlated with PC2. Furthermore, the first and second principal components (PC1 and PC2) explain nearly 75% of the variance of the data (Figure 3b), supporting the idea that the GC content of the codon’s third position of a gene is one of the most crucial factors for the robustness under mutational processes in human.

**Figure 3.**
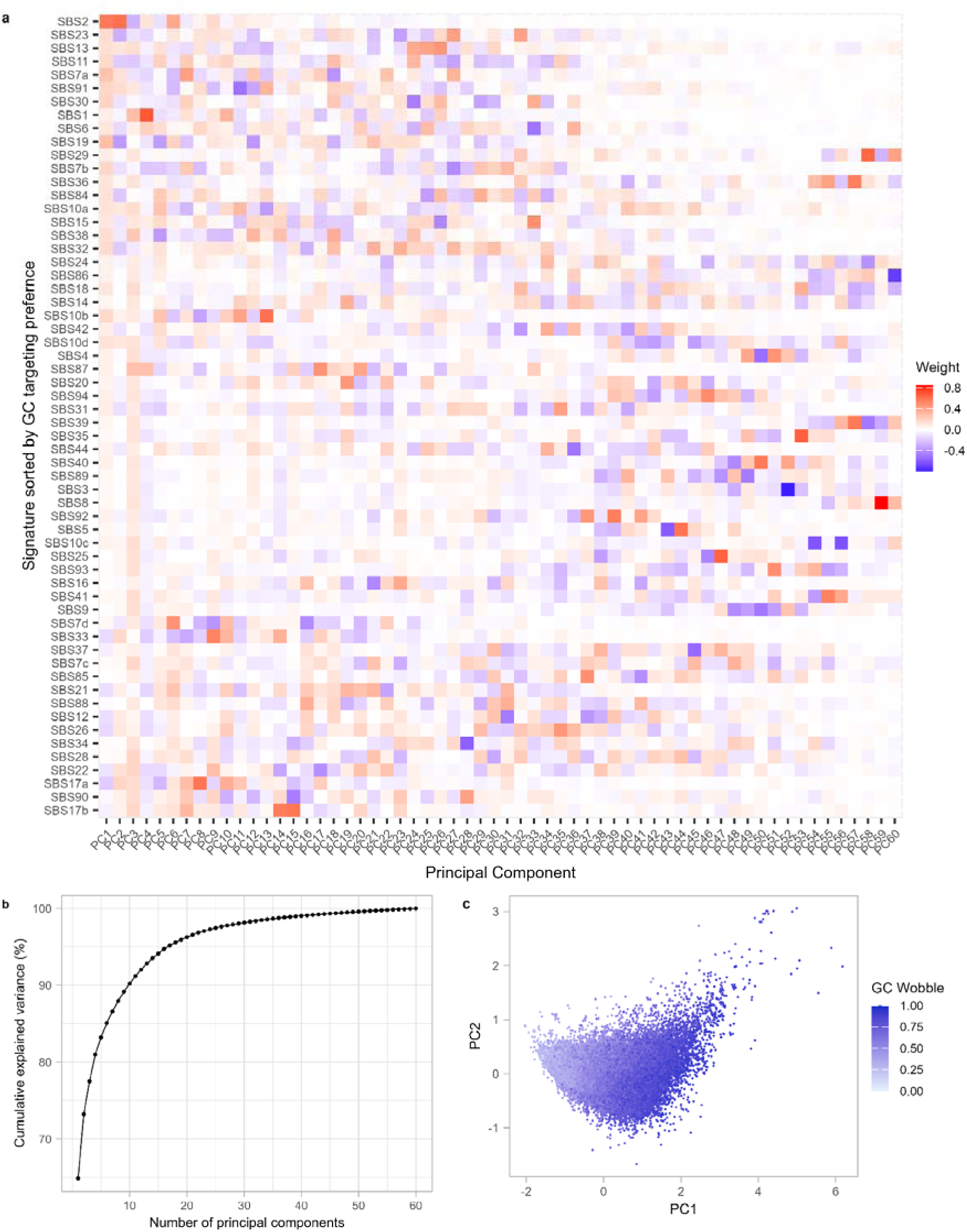
Principal component analysis (PCA) of mutational signature. **a.** Heat map colored by the weight of mutational signatures in each principal component. The Y-axis is sorted based on the GC targeting preference of each mutational signature. **b.** Shoulder plot of cumulative explained variance ratio. PC1 and PC2 explain 73% of the total variance, and from PC1 to PC10 explain 90% of the total variance. **c.** PC1-PC2 plot of all protein-coding sequences of the human genome colored by the GC contents at the wobble position. A sequence with high GC contents at the wobble position has a high PC1 value.

### The human genome is more robust under transition mutations than transversion mutations

We further investigated the weights of principal components from PC3 to PC10 to understand the variance remaining in the robustness profile of the human genome. The cumulative explained variance up to PC10 was 90%. We first focused on the mutational signature with the highest weight value in each principal component. SBS1 appeared to be the highest for PC3, PC4, and PC10; SBS10a for PC5; SBS7d for PC6; SBS7a for PC7; SBS84 for PC8; SBS22 for PC9 (Figure 4). These mutational signatures had different target preferences but commonly showed high specificity (Figure S6). To determine how the specificity of mutational signatures affects the robustness, we computed the specificity of mutational signatures using Shannon entropy. We found an arrow-tip shape from the average robustness under the mutational signature versus the specificity of the mutational signature plot (Figure 5a) and a negative correlation between the standard deviation of robustness and the specificity of the mutational signature (Figure 5b). As in the previous analysis, we assumed that GC targeting preference might explain the result. However, when we colored dots with the GC targeting preference, it appeared to have mixed results, although, as expected, there was a mild positive correlation along the y-axis. From the previous result, the human genome showed high robustness under C-to-T targeting mutational signature and low robustness under C-to-A targeting mutational signature. We further analyzed the robustness versus specificity plot by coloring the preference for the transition mutation, which makes pyrimidine-to-pyrimidine or purine-to-purine modification (Figure 5c). The color gradients clearly divided the arrow tip shape into two linear shapes intersecting where the mutational signatures induce almost random mutations. The linear regression on robustness versus transition mutation preference also showed a significant positive correlation (Figure 5d). These results showed the human genome is more robust under transition than transversion mutations.

**Figure 4.**
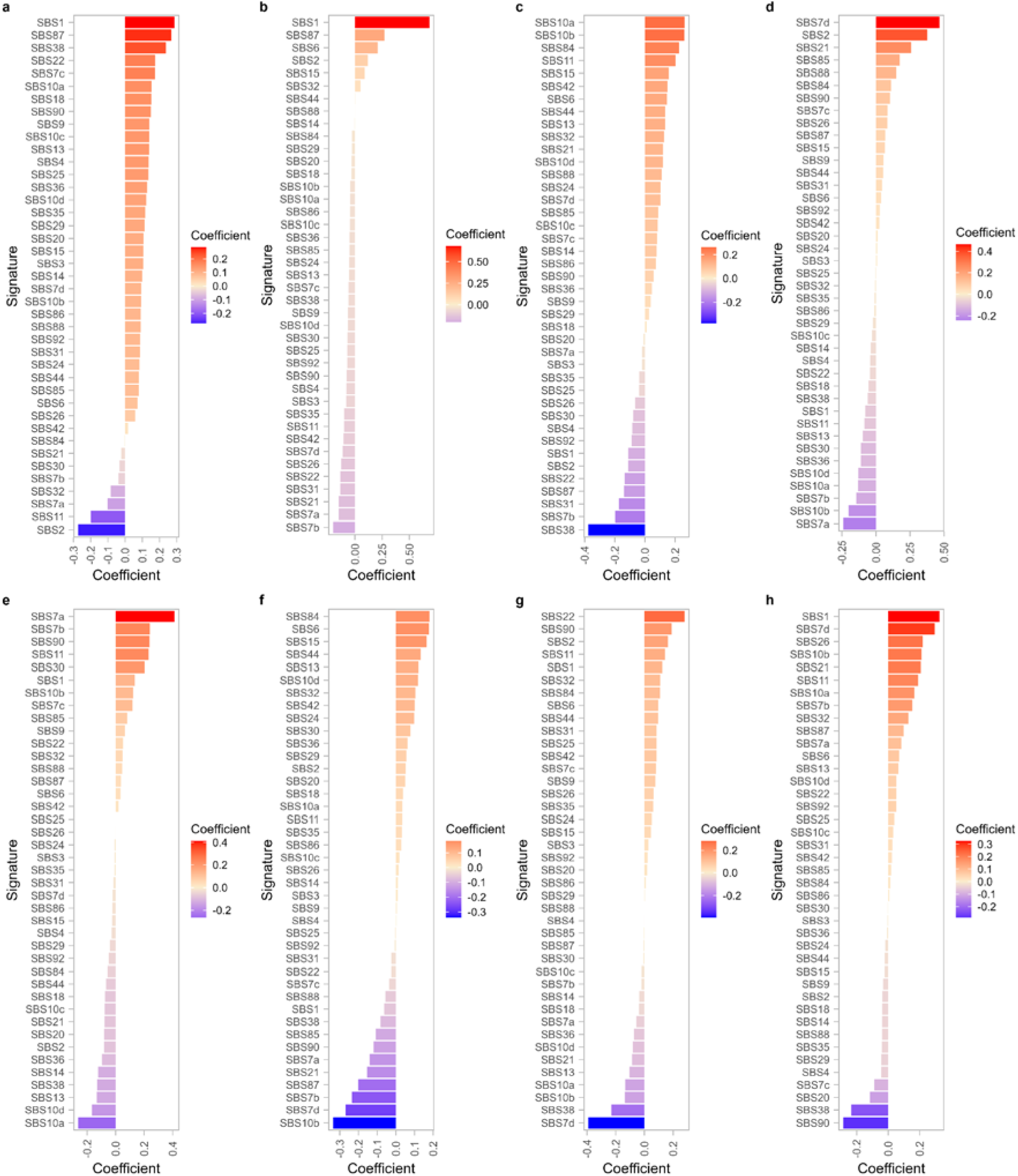
The weight of each mutational signature on PCs 1-10. The Y-axis is sorted based on the weight of each mutational signature of each principal component. SBS1 appeared to be the highest for PC3, PC4, and PC10 (**a, b,** and **h**); SBS10a for PC5 (**c**); SBS7d for PC6 (**d**); SBS7a for PC7 (**e**); SBS84 for PC8 (**f**); SBS22 for PC9 (**g**).

**Figure 5.**
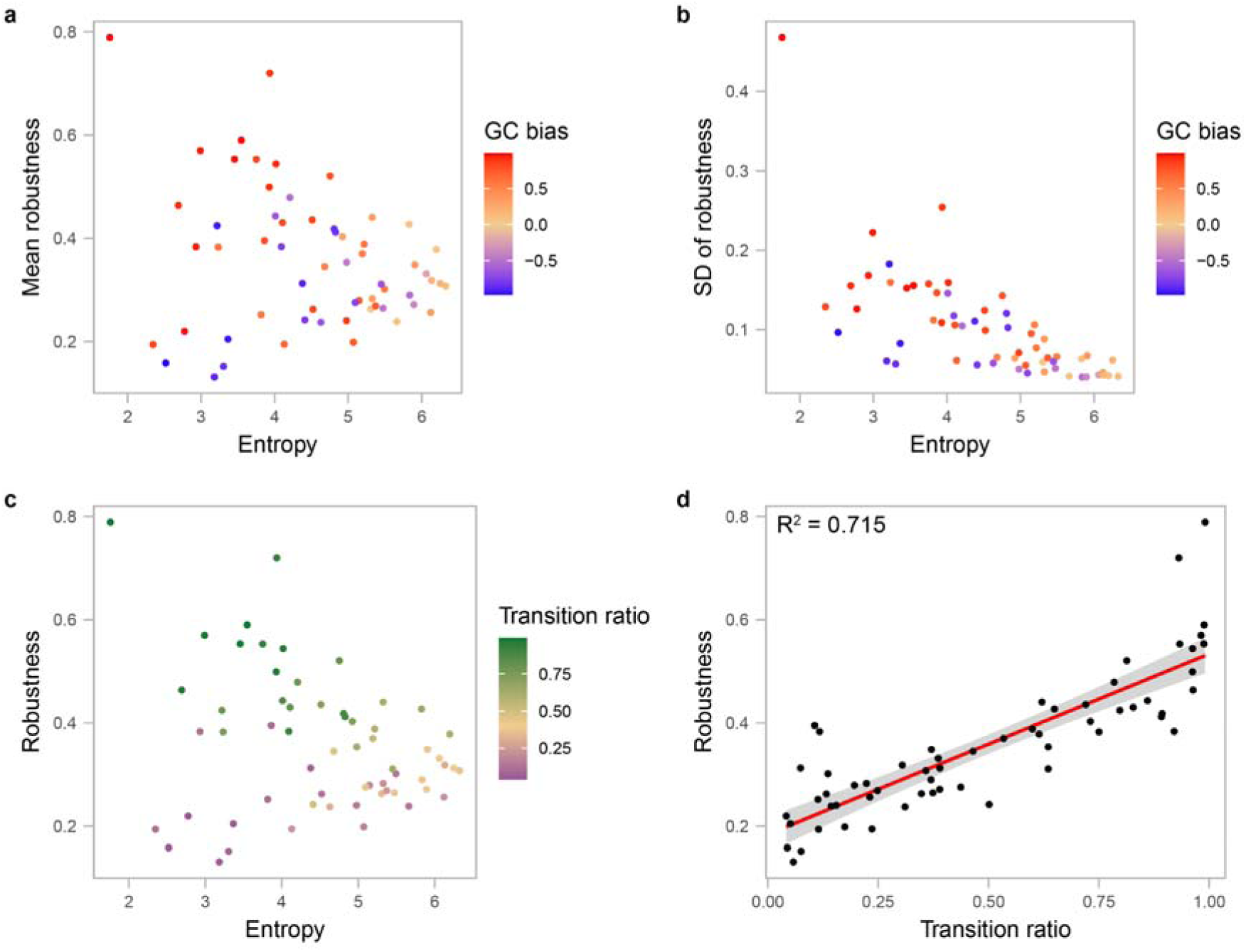
Impact of mutational signatures’ specificity and the transition mutation ratio on robustness. Specificity decreases with increasing entropy. **a.** The average robustness under mutational signature versus the specificity of the mutational signatures plot is colored by the GC targeting preference of the mutational signatures. **b.** The standard deviation of robustness under mutational signatures versus the specificity of the mutational signatures plot is colored by the GC targeting preference of the mutational signatures. **c.** Average robustness under mutational signature versus the specificity of the mutational signatures plot is colored by the transition mutation ratio of the mutational signatures. **d.** Linear regression plot of the average robustness of mutational signatures against the transition ratio of the mutational signatures. The two variables have a significant positive correlation (R^2^ = 0.715, F(1, 58) = 145.5, p < 2.2 x 10^-16^).

### Long-lived bat’s genome has higher robustness than the human genome under their native APOBEC3like enzymes activity

Several bat species have extremely high longevity and cancer resilience compared to mammals of similar size^22,23^. Bats also tolerate a diverse suite of viruses, seemingly asymptomatically, meaning they have evolved to control (some) viral infections without adverse inflammation^24^. One such mechanism involves the AID/APOBEC family of deaminase, which induces mutations in the viral genome, reducing viral fitness. We hypothesize that long-lived bats have higher robustness under APOBEC activity since they have had to avoid damaging their genome while maintaining high activity to suppress viral infection. To test our hypothesis, we measure motif specificity of bat APOBECs in cytidine deamination. An artificial DNA substrate of 369 bp was used to test the cytidine deamination activities of 6 bat APOBEC enzymes from the two species of Molossus molossus (short-lived bat) and Myotis myotis (long-lived bat). The substrate was designed to have multiple copies of each of the 16 NNC motifs that were randomly distributed in the sequence. This would provide relatively equal accessibility to each different NNC for the ABOBEC enzymes. The substrate DNA was incubated with each bat APOBEC enzymes, and then PCR amplified and followed by Next Generation Sequencing analysis. Figure 6 shows the average percentages of C to T mutation rates of all the same NNC in the full DNA sequence. As shown in the figure, each of the four APOBEC3A-like and APOBEC3C-like enzymes showed distinctively different motif preference patterns. In contrast, the APOBEC1-like enzymes from the two species had the same preference patterns. While each of the 6 enzymes worked similarly well on their different sets of multiple NNCs, it was interesting that all of them had very low activities on motifs such as GGC and AGC. We then evaluated the robustness of two bat genomes, *Molossus molossus* and *Myotis myotis*, under 6 COSMIC mutational signatures, and the activity of APOBEC1, APOBEC3A, APOBEC3C from these bat species using Sinabro (Figure 7). We found that *M. myotis* has higher robustness under APOBEC activity, especially under SBS2, than humans and *Mol. molossus* (Figure 6e and 6f; Mann–Whitney U test p < 2.2x10^-16^). This trend was evident when we compared robustness under their native APOBEC3A (Figure 6g). We used SBS4 (tobacco smoking) signatures as negative control and, as expected, did not show a difference (Figure 6b). Although we expected to observe higher robustness to other mutational signatures in the long-lived bat, we found no or small differences in SBS1, SBS4, and SBS6 (Figures 6a, 6c, and 6d). This suggests that bat longevity and cancer resilience might have cross-evolved under their own APOBEC activity but not under other documented mutagens to humans.

**Figure 6.**
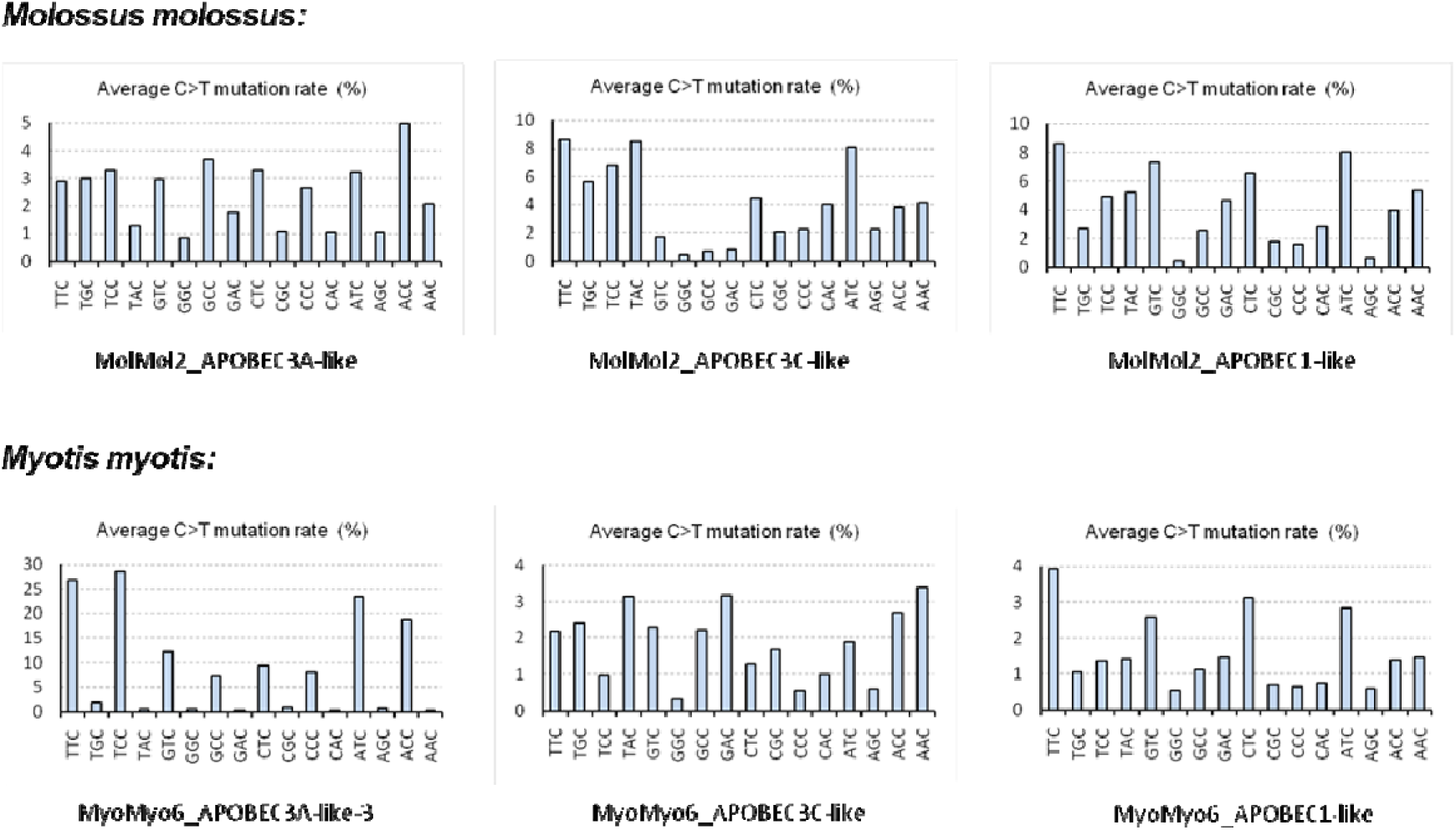
Motif preference patterns of bat APOBECs in cytidine deamination. Each bat APOBEC enzyme was incubated with a 369 bp artificial DNA substrate, followed by PCR amplification of the resultant DNA and Next Generation Sequencing analysis. Each bar shows the average percentage of the C to T mutation rates of the corresponding NNC from the full length of the DNA sequence.

**Figure 7.**
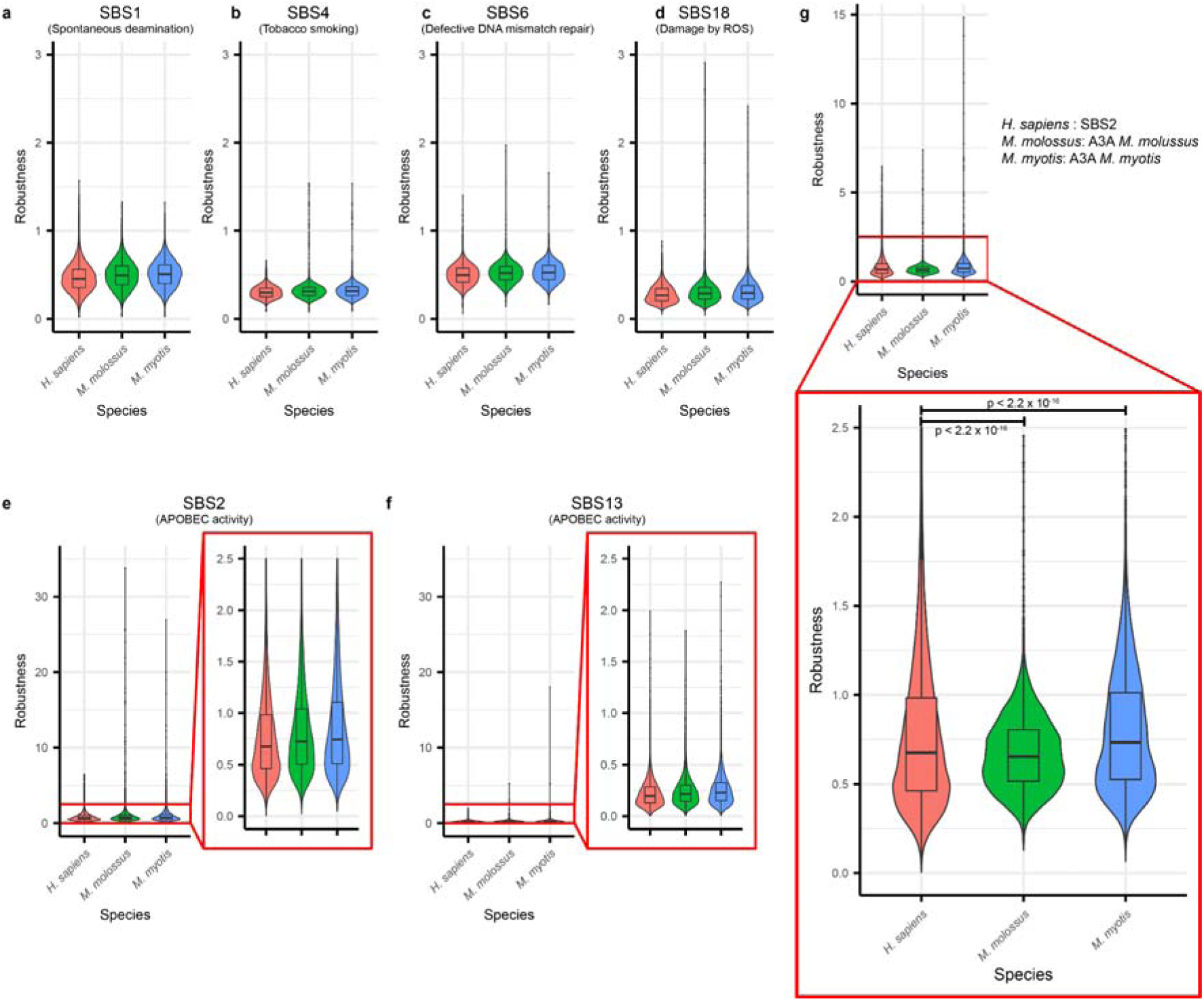
Comparison of robustness under mutational signatures between human and two bat genomes. Robustness under COSMIC mutational signature **(a)** SBS1 spontaneous deamination, **(b)** SBS4 tobacco smoking, **(c)** SBS6 defective DNA mismatch repair, **(d)** SBS18 damage by ROS, **(e** and **f)** SBS2 and SBS13 APOBEC activity. Pair wise Wilcoxon rank sum test in Table S2. **g.** Robustness under their native APOBEC3A activity. *M. myotis* showed the highest and *M. molossus* showed the lowest robustness.

## Discussion

Genomes are constantly exposed to various mutational drivers, from spontaneous deamination to specific mutagens, such as UV exposure. Tolerance to these mutational burdens is crucial to maintaining a robust biological system. Ironically, cells are both evolvable and robust to the mutational stressors in their microenvironment. In a 2008 paper ^6^, Wagner introduced genotype robustness as the number (or fraction) of neutral neighbors of the genotype. Greenbury et al. expanded Wagner’s definition, allowing more than one mutation and introducing the concept of n-robustness ^7^. However, these definitions are too broad to evaluate the robustness of a gene directly under specific mutational drivers. To resolve this problem, we defined a gene’s robustness under a specific mutational process using sequential mutations under a mutational signature.

Here, we propose a simple method for computing the robustness under a specific mutational process for a given coding sequence, Sinabro, to simulate single base substitution under a given mutational driver. Robustness was measured as the average number of single base substitutions a given sequence can tolerate. Our simulator successfully recapitulated COSMIC mutational signatures. Our results highlight that the human genome displays variable robustness across different mutational signatures, with notable resilience under the APOBEC mutational signature, SBS2. This signature is linked to a cytidine deaminase involved in viral defense mechanisms, but it has also been implicated in cancer pathogenesis and tumor heterogeneity. As many research studies highlight APOBEC mutagenesis as part of carcinogenesis and cancer initiation, robustness to SBS2 is a key finding. Our findings validate our previous discovery on the role of APOBECs in later stages of cancer by contributing to the heterogeneity of cancer cells that are more evolvable and probably less robust to mutations^25^.

We also demonstrate the utility of our approach when comparing robustness to mutagenic processes across species. Among bats, *M. myotis* (maximum lifespan = 37 years) and *Mol. molossus* (maximum lifespan = 5.6 years) represent extremes in the relationship between lifespan and body mass^26^. While the expansion of the APOBEC3 gene subfamily in bats has been well documented^27,28^, ours is the first analysis to explore the implications for genomes of more than one bat species. Our finding of varying robustness to their own (Fig. 6g), but not other species’ APOBEC3 activity suggests coevolution between the genome and native APOBEC3. This contrasts with our previous finding of similar APOBEC3 motif representation across cancer and non-cancer genes in *Pteropus alecto* (maximum lifespan = 20.3 years)^25^, a bat with the most APOBEC3 catalytic domains known to date, 13^28^. But, consistent with the coevolution hypothesis, we also found distinct distributions in inferred mutational susceptibility to APOBEC3-mediated deamination^25^. Our focus on robustness illuminates another dimension of genome variability; while the long-lived bat displays higher robustness, the short-lived bat shows lower robustness and its frequency distribution is distinct from those of both human and the long-lived bat. Thus, our finding that APOBEC3 mutational signatures strongly influence the robustness of the human genome is validated across bats and appears to relate to the evolution of extreme lifespans as well.

## Methods

### Sinabro: sequential mutation simulator

The main function of Sinabro is to generate sequences by simulating a single base substitution at a time. To accomplish this function, Sinabro applies the custom Python class Trajectory, which stores the input sequence and other sequences generated by sequential mutation. The class also stores mutations in HGVS mRNA format, HGVS Protein format, and mutation type. The mutation type in this study is a string formatted as (5’ context)[(original nucleotide)>(mutated nucleotide)](3’ context) to record the context and the type of single base substitution. For example, a mutation type T[C>T]A represents the C-to-T mutation in TCA. Users can either manually create a Trajectory class object by inserting records or automatically fill it using predefined methods with parameters (details in Supplementary Note S2). To compute the robustness of a given coding sequence under mutational signatures, we used parameters condition=“nonsynonymous” and method=“signature.” With these parameters, Sinabro reads a signature file containing the occurrence frequencies of 96 mutation types and converts it to a 4-by-64 matrix where columns are all possible 64 trinucleotides, and rows are the nucleotides resulting from mutation. For example, f_2,1_ represents that the center nucleotide is mutated to nucleotide 2 (C) in the context of trinucleotide 1 (AAA). Using the matrix, Sinabro then computes a 4-by-n probability matrix whose element p_i,j_ is the probability of j-th nucleotide in the sequence mutated to i-th nucleotide, where n is the length of a given sequence. Then, a single base substitution is selected based on the probability matrix, and the resulting sequence, HGVS mRNA format, HGVS Protein format, and mutation type are recorded to the Trajectory object. Sinabro decides whether to continue the process by checking whether the resulting mutation changed the codon. Sinabro ends the automatic filling of the Trajectory object if the mutation is non-synonymous. To compute the robustness under bat’s APOBEC activity, we used parameters condition=“nonsynonymous” and method =“mut_types.” With these parameters, Sinabro detects all potential mutation sites and computes the probability of mutation of each site based on the given probabilities of each mutation type. Then, the same procedures, selecting a single base substitution and checking codon difference, are repeated to measure robustness.

We validated that Sinabro faithfully recapitulates mutational signatures by simulating a single base substitution on 10,000 per random sequence of 2,200 bp long. Then, we re-created each mutational signature based on the simulation result (Figure S1), and the cosine similarity between the original mutational signatures was computed (Figure S2).

### Data and preprocessing

We downloaded the FASTA file containing all protein-coding transcript sequences from GENCODE (Release 40). Then, only the complete protein-coding region of each record was extracted. In detail, we extracted substrings of the record containing coding sequence using the FASTA file name field annotation. Those sequences were filtered to have the correct length of a multiple of three, start codon, and stop codon. In addition, we removed sequences from the pseudoautosomal region of the Y chromosome. For the computation of the robustness under mutational signature, each sequence also includes two additional nucleotides, one at the 5’ end and one at the 3’ end. Those nucleotides were required to simulate mutations at the first and the last position in trinucleotide contexts of COSMIC mutational signatures. The resulting FASTA file with a total of 73,214 coding sequences was used in further robustness computation.

### Computation of the robustness under mutational signatures

We defined the robustness of a coding sequence under the mutational signature as the average number of synonymous single-base substitutions under the mutational signature. We wrote a custom Python script computing the robustness of a single sequence utilizing Sinabro. The script generates 1,000 Trajectory objects per mutational signature from a coding sequence record in the preprocessed GENCODE file. The script then makes a 1,000-by-79 Pandas DataFrame that stores l-1, where l is the length of the Trajectory object and saves it into a CSV file. We used the SGE high-performance computing cluster in the Laufer center to compute the robustness for all coding sequences in the processed GENCODE file by submitting an array job. We calculated the robustness of each coding sequence by averaging the results from each output file and combined them into a single file, all_gencode_rums_profile.csv, which was used for further analysis. The robustness profile file was imported into R for the ranking plots, and violin plots of the robustness distribution were created using the ggplot2 package^29^ (Figure 2).

### Calculation of GC targeting preferences and entropy of mutational signatures

COSMIC mutational signatures were downloaded from COSMIC. GC targeting preferences of each mutational signature were computed by summing the percentage of C>A, C>G, and C>T. The entropy of the mutational signature was calculated as H(s)=-Σp(s_i_)log_2_p(s_i_) where s is a mutational signature and p(s_i_) is the percentage of mutation type i. We used Numpy to compute GC targeting preferences and entropy, and the results were exported to a single CSV file.

### GC contents at the wobble position of coding sequences

GC contents at the wobble position, that is, the fraction of G or C at the wobble positions in the given coding sequence, were computed using a custom Python script. The script reads in the preprocessed GENCODE FASTA file with the BioPython SeqIO module. The GC content at the wobble position is calculated by dividing the number of G or C at the wobble position by the number of wobble positions, which is the total length divided by three. The result was then exported to a single CSV file.

### Transition mutation ratio of mutational signatures

The transition mutation ratios of mutational signatures were computed by summing the percentage of C>T and T>C mutations for each mutational signature using Numpy and exported to a single CSV file.

### Principal component analysis

COSMIC mutational signatures include signatures likely to arise from sequencing artifacts. Hence, we excluded 19 possible sequencing artifact signatures and performed principal component analysis (PCA). We used the “PCA.fit()” function from “sklearn.decomposition” with default parameters to perform the principal component analysis on the subsetted robustness profiles matrix. Then, the cumulative explained variance ratio and the weights of principal components were exported into a single CSV file using the Numpy cumsum() function on PCA.fit.explained_variance_ratio_ and PCA.fit.components_, respectively.

All generated files were imported into R, and the principal components heat map, cumulative explained variance plot, PC1-PC2 with GC contents of coding sequences, principal component weight plots, robustness versus entropy plots, and robustness to transition ration regression plot were created using the ggplot2 package.

### Bat APOBEC protein expression, purification and cytidine deamination motif preference analysis

The bat APOBEC gene coding sequences of Molossus molossus and Myotis myotis were identified in the bat1k longevity project. The genes were cloned into a pPICZ yeast expression vector for expression and purification in yeast as previously described^30^. For purification purposes, a GST-tag was introduced in frame at the 5’-end of each gene so that a fusion protein would be expressed. Screening of transformants and expression of GST-APOBEC fusion proteins were carried out as previously described^30,31^. Cells were lysed using a French Pressure Cell Press. GST-tagged bat APOBEC proteins were purified using Glutathione Sepharose High-Performance beads (GE Healthcare, Cat # 17-5279-01) as previously described^30^. Purified proteins were stored in a solution containing 20 mM Tris-HCl pH 7.5, 100 mM NaCl, 1 mM DTT and 5% glycerol.

To examine the cytidine deamination (C to T mutation) activities of the purified bat APOBEC enzymes, we employed a PCR-based assay described previously ^31^ where in Next Generation Sequencing (NGS) was used to amplify and sequence a DNA substrate incubated with the APOBECs. The substrate DNA fragment was a 369 bp-long artificial sequence containing multiple copies of all the 16 NNC motifs and cloned into the pcDNA3.1 vector. This DNA fragment was flanked at both the 5’- and 3’-ends with AT-only sequences to ensure unbiased PCR amplification post APOBEC-mediated cytidine deamination. The purified bat APOBEC enzymes were incubated with the plasmid DNA substrate at 33 °C, pH6.5 for 1 h, and then PCR amplified using Taq DNA polymerase and primers annealing to the AT-only flanking regions. PCR products were purified and NGS was performed by GENEWIZ (Azenta Life Sciences). The sequencing data were analyzed at the Galaxy web platform (usegalaxy.org) using tools FastQC, Trimmomatic, Bowtie2, Naïve Variant caller and Variant Annotator. Output data were analyzed for NNC hotspot motif preference of the bat APOBECs by comparing the normalized frequency of mutations found at each motif.

### Computation of probability of mutation type for measuring robustness under bat APOBECs

To measure robustness using Sinabro, the conditional probability that the 5’ context is one of NNC, given that a C-to-T mutation occurs. In detail, denote the probability of a single C-to-T mutation as P(A) and the probability that the 5’ context of mutated C is c_i_ as P(B_i_) where c_i_ is an element of a set of all possible 5’ context C={c_i_}. Then, P(B_i_|A)=m_i_/M where mi is the number of C-to-T mutations in the context ci, and M is the total number of C-to-T mutations. These probabilities were computed for each APOBECs and passed to the parameter mut_type_prob in Sinabro for measuring robustness along with other parameters condition=“nonsynonymous” and method =“mut_types.”

## Data availability

All human protein-coding transcript sequences are available in GENCODE (release v40, https://www.gencodegenes.org/human/release_40.html). Genome assembly and mRNA annotation of two bat species, *M. molossus* (molMol2) and *M. myotis* (myoMyo6), are available in Bat1K genomes from Hiller Lab (https://bds.mpi-cbg.de/hillerlab/Bat1KPilotProject/).

## Code availability

The sequential mutation simulator Sinabro is available on the Sinabro GitHub repository (https://github.com/StudyingAnt/sinabro). All computational analysis scripts are available on the GitHub repository (https://github.com/StudyingAnt/Simple_robustness_analysis_of_cds).

## Acknowledgments

We gratefully acknowledge funding from the Physical Sciences Oncology Network at the National Cancer Institute (grant U01CA261841) as well as support from the Stony Brook Cancer Center.

This work was also supported partly by R01 grant R01CA272601 (NCI). L.M.D was supported in part by NSF-IOS 2032063 and NSF-DEB 1838273.

## Supplementary materials

### Incorporation of mutational processes into the robustness

Wagner straightforwardly defined the robustness of genotype g as the number (R_g_) or fraction (r_g_) of neutral genotypes separated by 1 nucleotide (Figure S7A). If the genotype length is L, there are 3L genotypes separated by 1 nucleotide considering only substitutions hence, we have a relation r_g_=R_g_/3L. We can view r_g_ from a different perspective. One can think of r_g_ as an expectation that the genotype maintains its phenotype after a random single nucleotide substitution where all mutation types have the same probability of 1/3L. Applying a similar perspective, the n-robustness of genotype g (r_g_^(n)^) can be considered as an expectation that the genotype maintains its phenotype after n single nucleotide substitution where the probability to be mutated into a specific genotype is the same for all genotypes. One can immediately think of considering different probabilities of being mutated into different genotypes. Then, the robustness of genotype g under mutational process can be written as $$r_g=\sum_{g_i \in G}{P(g>g_i) }$$ where P(g>g_i) is a probability of g mutated to g_i, and G is a set of all genotypes having the same phenotype with genotype g separated by single nucleotide substitution. Furthermore, one can consider sequential mutations until phenotype changes and the length of a path from g to g_i instead of restricting the number of mutations to a constant n. We can define robustness as the expectation of the length of a path. These can also be interpreted as the number of mutations given by the genotype that can be tolerated before phenotype changes. In this study, a protein-coding sequence and its variants by mutational signatures are the genotype g and g_i, respectively, and their translated amino acid sequences are phenotypes.

### Supplementary Note S2. Parameters of Sinbaro auto-filling function

Currently, in Sinabro, we have two stop conditions and three methods for mutation: max_length and nonsynonymous; random, mut_type, mut_types, and signature. The first stop condition, max_length, runs the simulation until the sequence cannot be mutated by the given mutation method or the length of the Trajectory reaches the given max_length. The second stop condition, nonsynonymous, runs the simulation until the sequence gets a non-synonymous mutation. The first mutation method, random, generates random single base substitution, meaning all nucleotides in the sequence have the same probability of mutation and the same probability of being mutated to one of three other nucleotides. The second mutation method, mut_type, runs the simulation of a given mutation type. It searches for candidates that match the given nucleotide context of the mutation type, randomly selects a position to mutate, and mutates the sequence based on the given mutation type. The third mutation method, mut_types, is similar to mut_type but used when each contexts have different probability to be mutated. It runs a simulation of a given mutation types and according probability of each mutation. The final mutation method, signature, runs a simulation of a given mutational signature as described in the method section. The mutational signature can be given by its COSMIC mutational signature name or custom mutational signature either generated by SigProfiler or manually created by the user.

**Figure S1.**
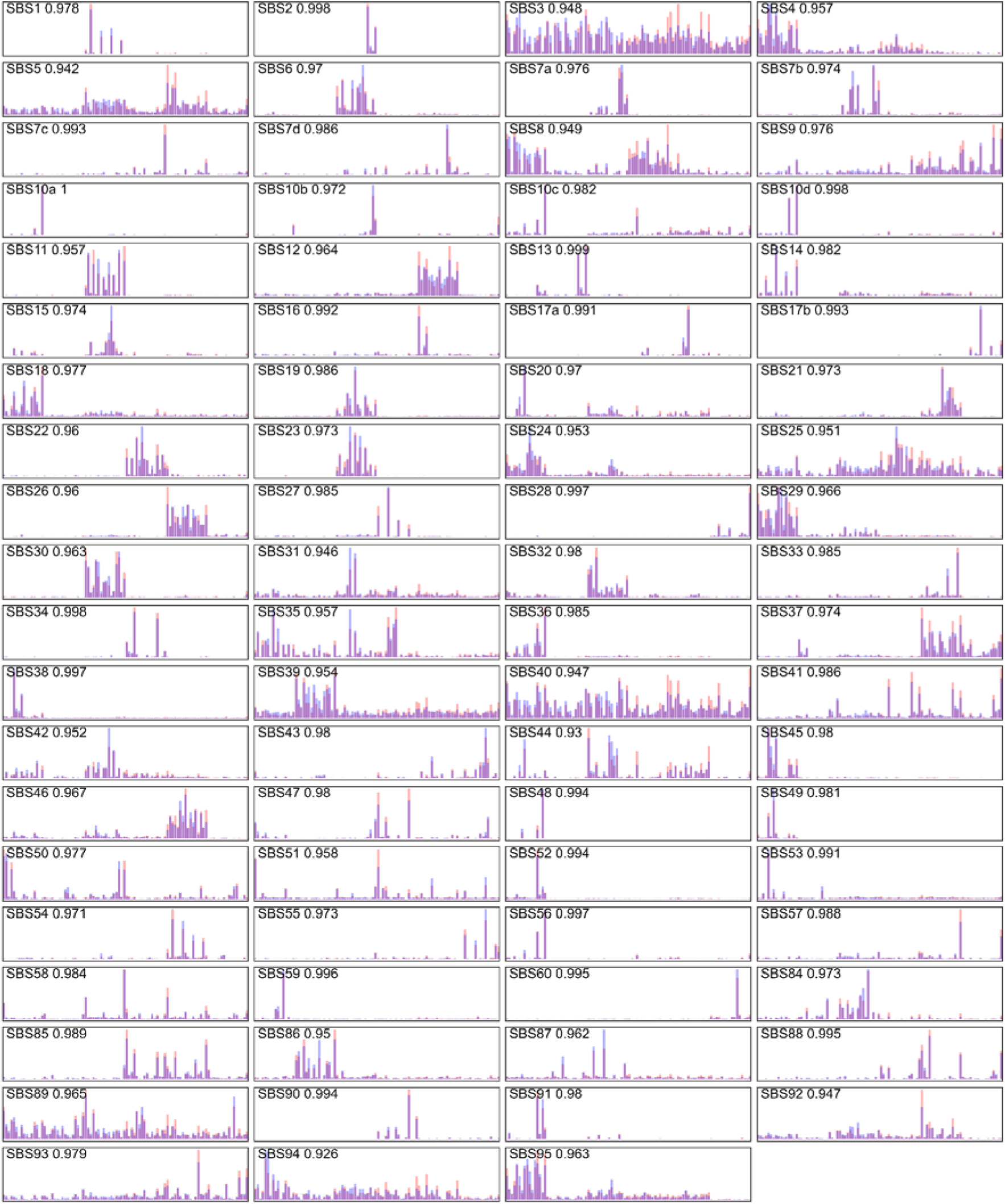
Recapitulation of all COSMIC mutational signatures using Sinabro. Randomly generated 10,000 of 2,200 bp long sequences with GC contents of 0.4 were passed to Sinabro, and simulated single mutations on those sequences to validate Sinabro faithfully recapitulates mutational signatures. The number in each plot is the cosine similarity between the original COSMIC mutational signature and simulation.

**Figure S2.**
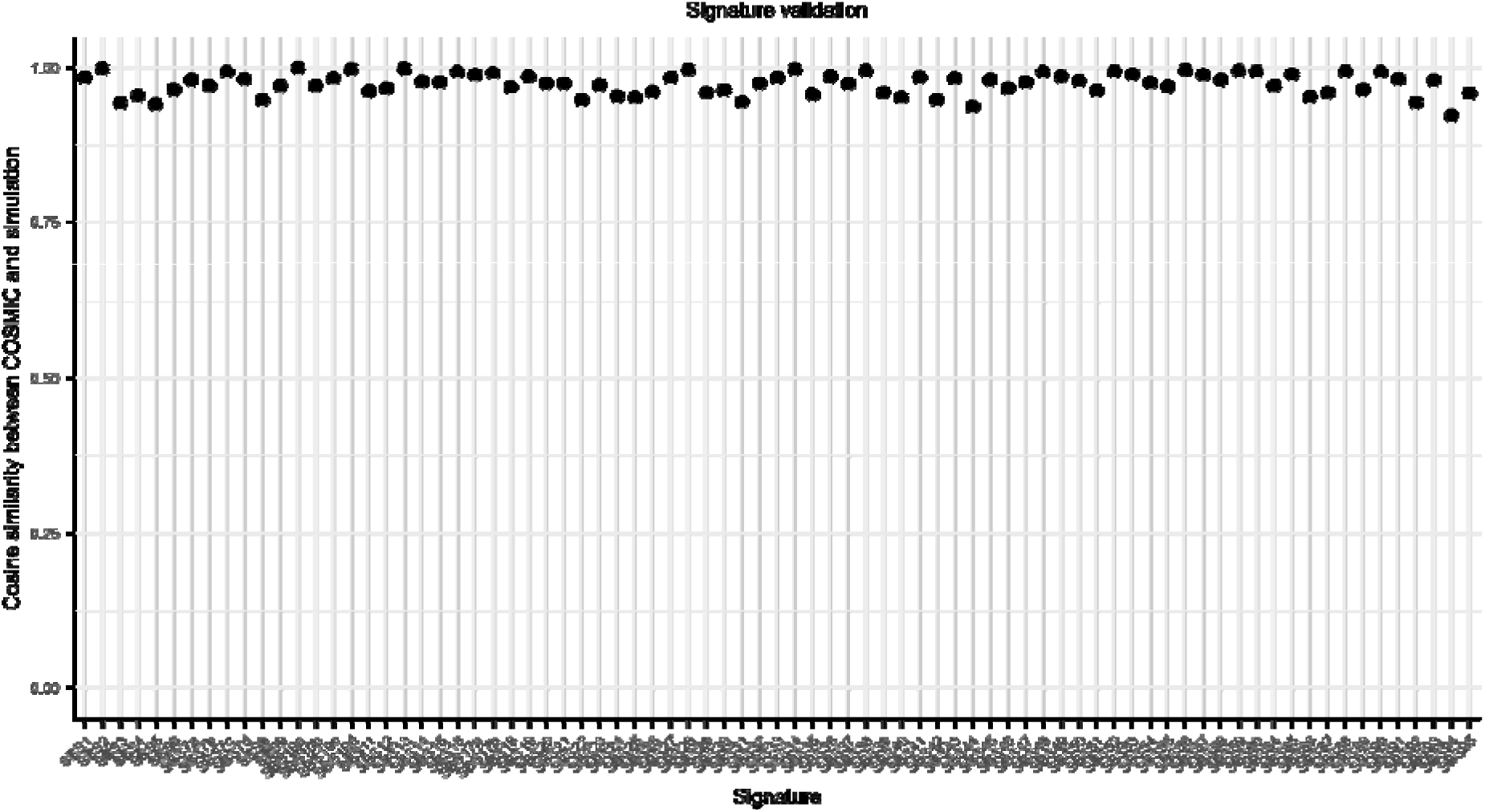
Cosine similarity between COSMIC mutational signature and Sinabro simulation. The average cosine similarity was 0.974, the maximum 1 for SBS13, and the minimum 0.926 for SBS94.

**Figure S3.**
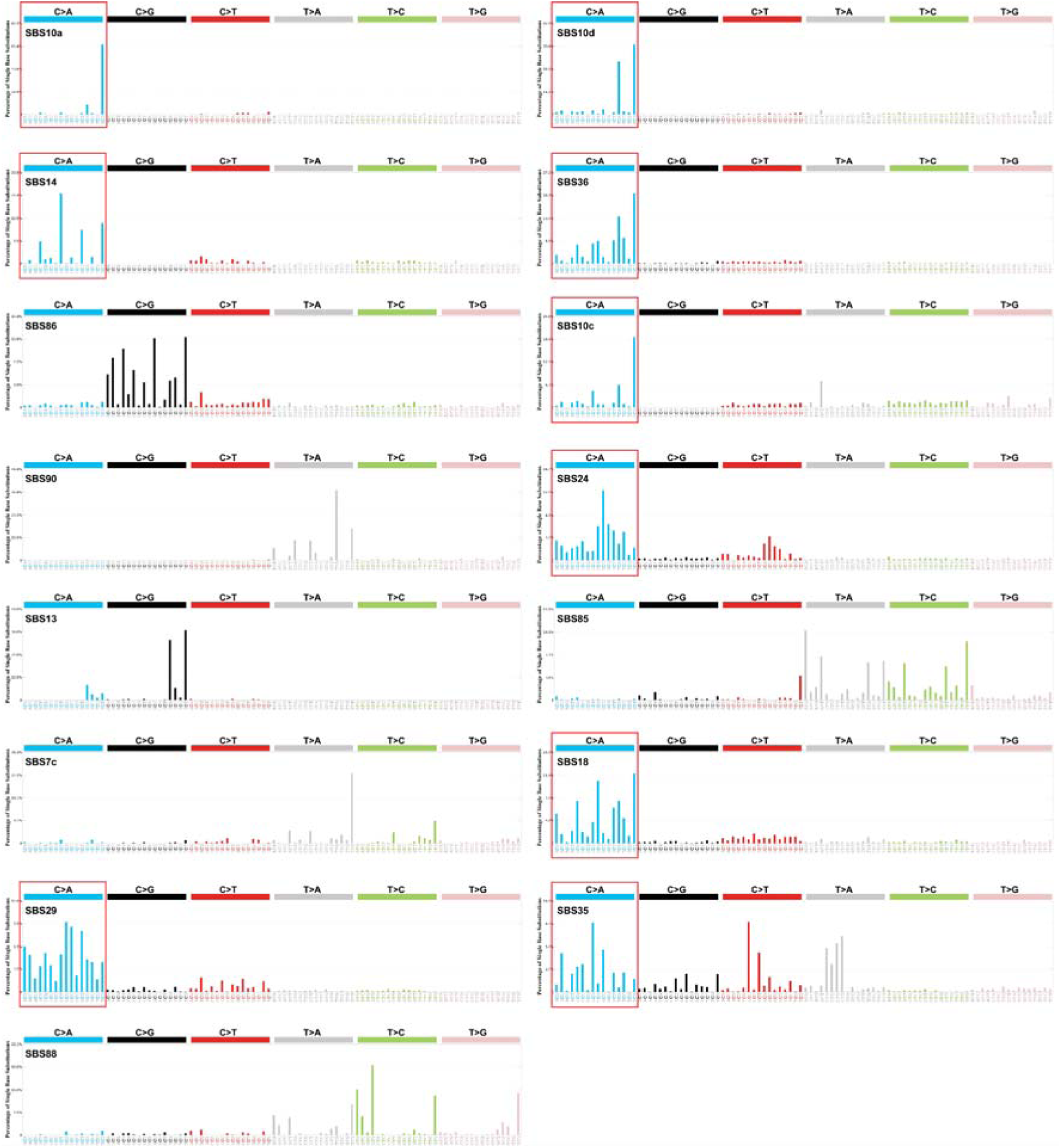
The bottom 15 mutational signatures ranked by the average robustness. 9 out of 15 signatures are almost exclusively inducing C-to-A mutation.

**Figure S4.**
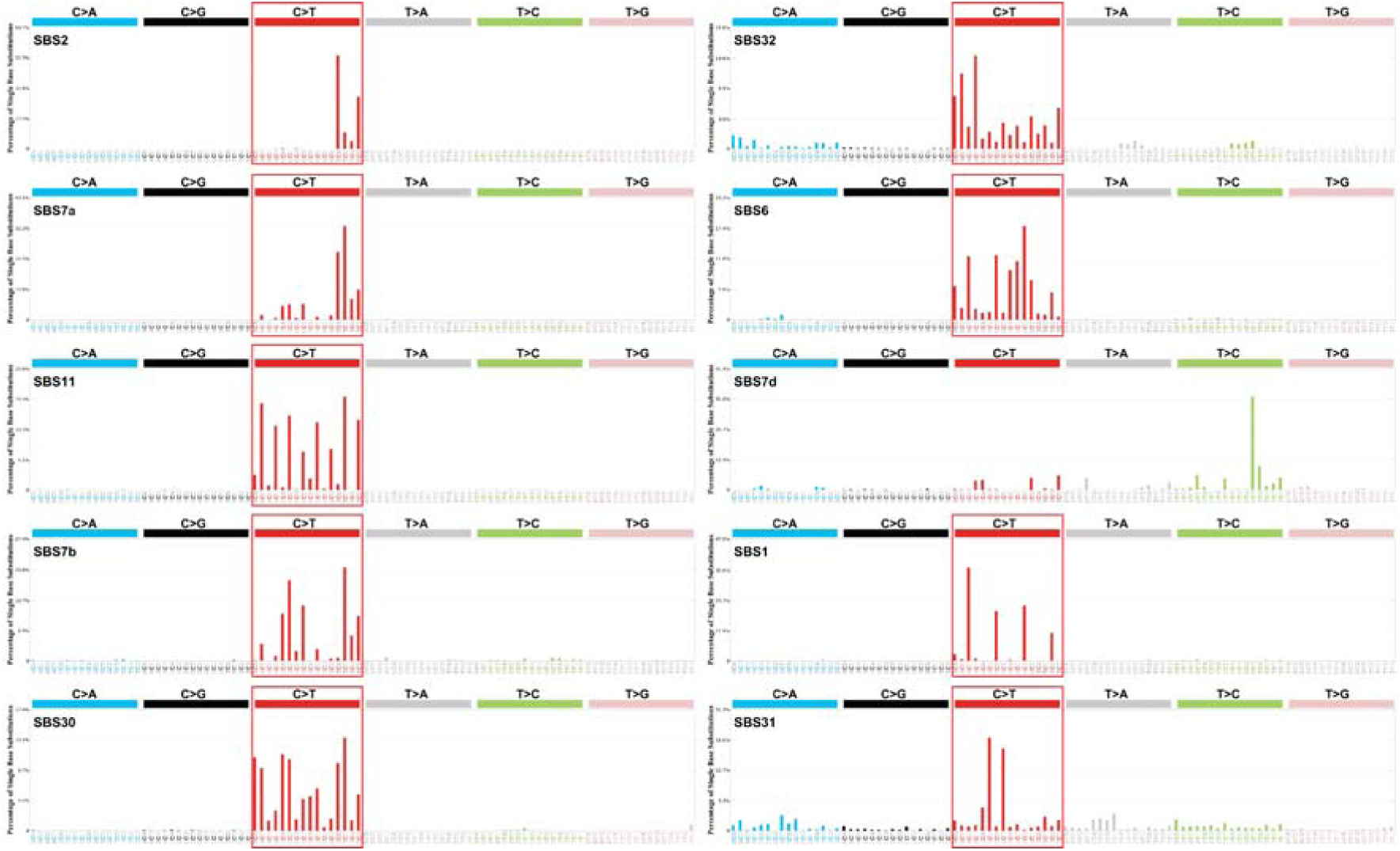
The top 10 mutational signatures ranked by the average robustness. 9 out of 10 signatures induce almost exclusively C-to-T mutation.

**Figure S5.**
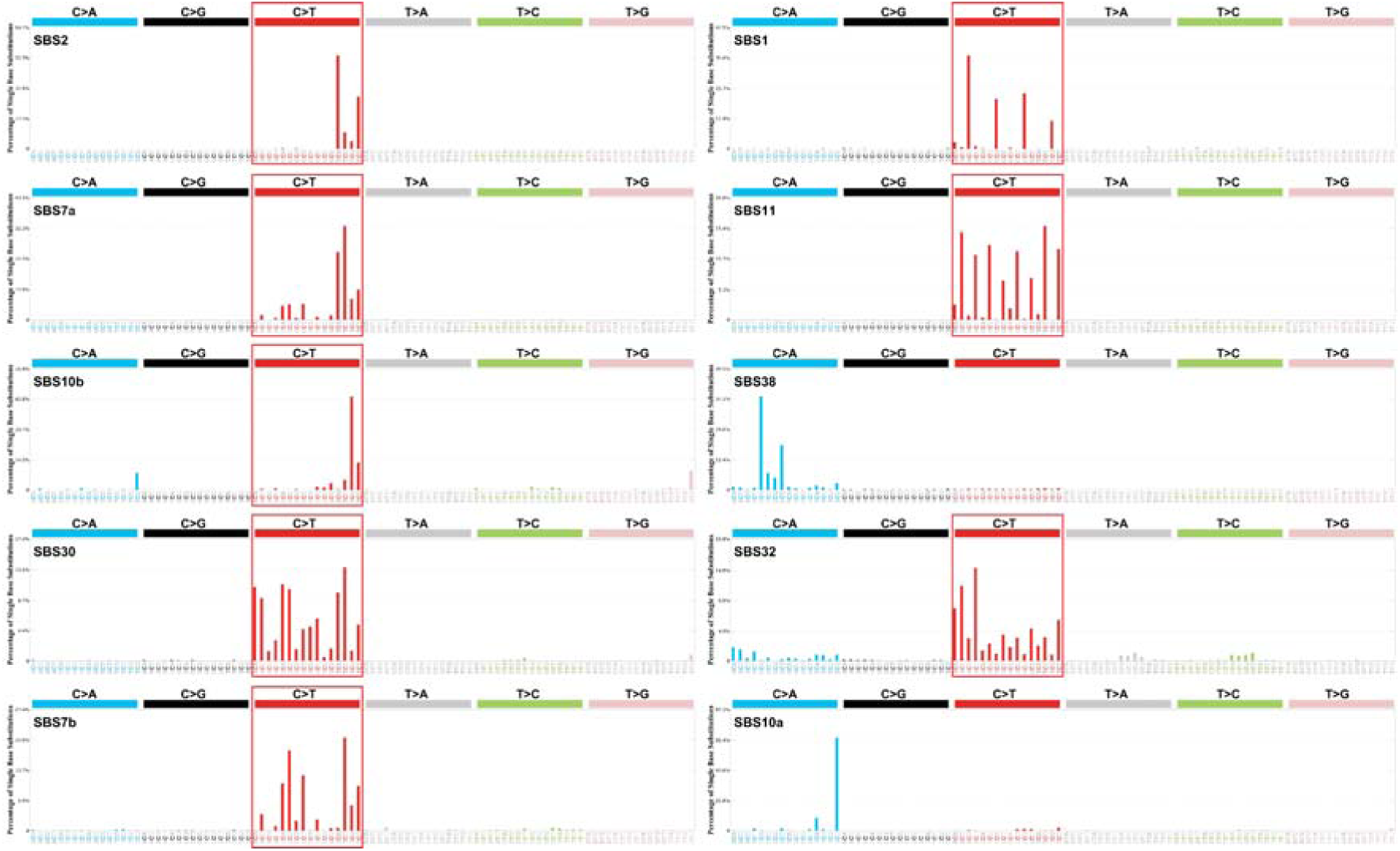
The top 10 mutational signatures ranked by the standard deviation of robustness distribution. 8 out of 10 signatures induce almost exclusively C-to-T mutation.

**Figure S6.**
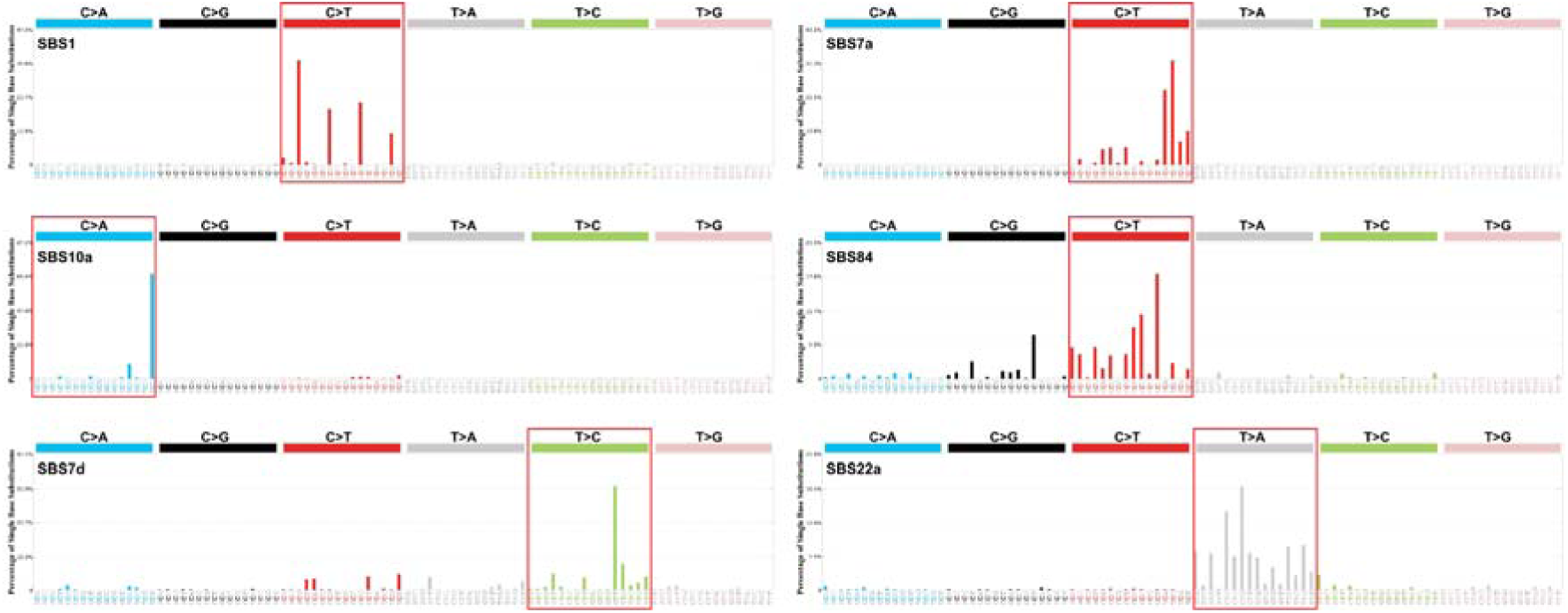
The mutational signatures with the highest weight of each principal components. Mutational signatures induce a specific mutation rather than a random mutation.

**Figure S7.**
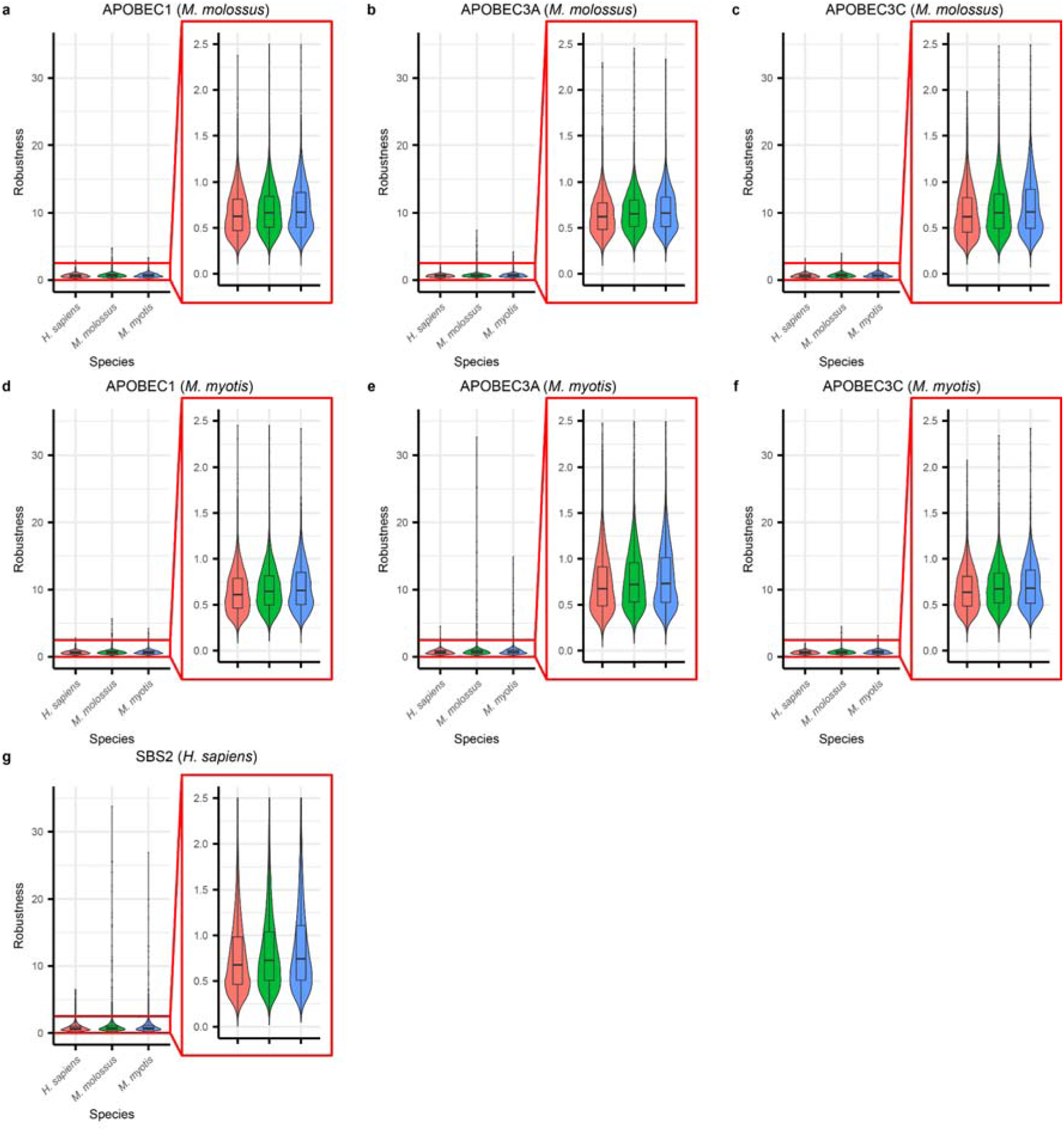
Robustness under the activity of APOBECs from different species. **a.**, **b.**, and **c.** Robustness under APOBEC1, APOBEC3A, and APOBEC3C of *M. molossus*, respectively. **d.**, **e.**, and **f.** Robustness under APOBEC1, APOBEC3A, and APOBEC3C of *M. myotis*, respectively. **g.** Robustness under SBS2 from *H. sapiens*. Pair wise Wilcoxon rank sum test in Table S3.

**Figure S8.**
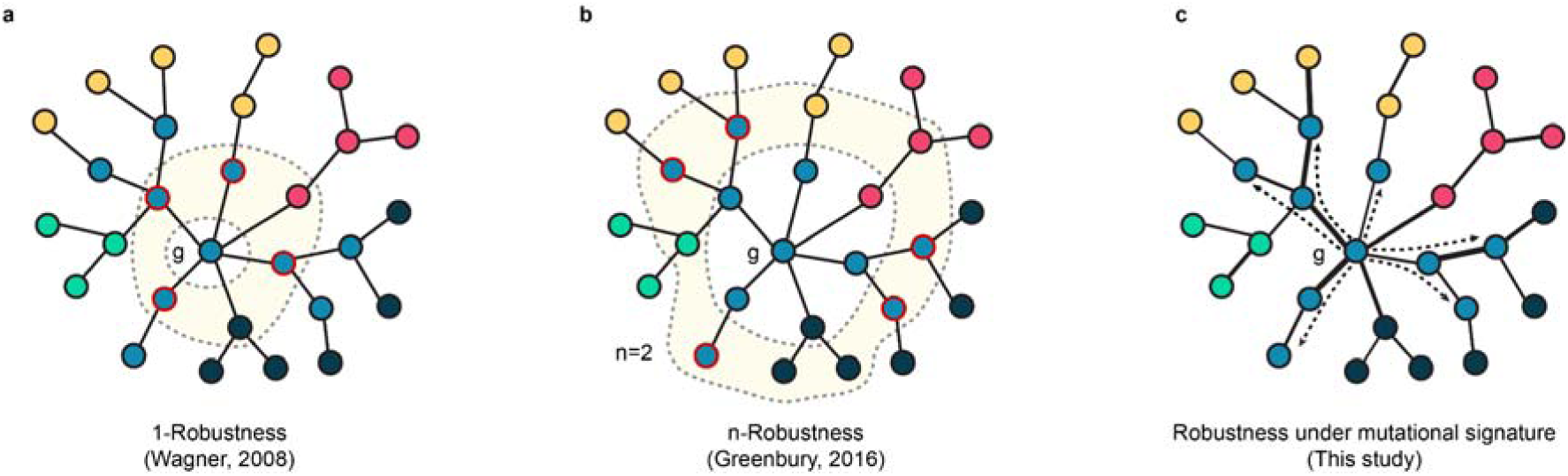
Schematic comparison between the definitions of robustness. **a.** 1-robustness definition by Wagner. **b.** n-robustness definition by Granbury. **c.** Robustness under mutational signature used in this study.

